# Variability in donor lung culture and relative humidity impact the stability of 2009 pandemic H1N1 influenza virus on nonporous surfaces

**DOI:** 10.1101/2023.04.17.537242

**Authors:** Zhihong Qian, Dylan H. Morris, Annika Avery, Karen A. Kormuth, Valerie Le Sage, Michael M. Myerburg, James O. Lloyd-Smith, Linsey C. Marr, Seema S. Lakdawala

## Abstract

Respiratory viruses can transmit by multiple modes, including contaminated surfaces, commonly referred to as fomites. Efficient fomite transmission requires that a virus remain infectious on a given surface material over a wide range of environmental conditions, including different relative humidities. Prior work examining the stability of influenza viruses on surfaces has relied upon virus grown in media or eggs, which does not mimic the composition of virus-containing droplets expelled from the human respiratory tract. In this study, we examined the stability of the 2009 pandemic H1N1 (H1N1pdm09) virus on a variety of nonporous surface materials at four different humidity conditions. Importantly, we used virus grown in primary human bronchial epithelial (HBE) cultures from different donors to recapitulate the physiological microenvironment of expelled viruses. We observed rapid inactivation of H1N1pdm09 on copper under all experimental conditions. In contrast to copper, viruses were stable on polystyrene plastic, stainless steel, aluminum, and glass, at multiple relative humidity conditions, but greater decay on ABS plastic was observed at short time points. However, the half-lives of viruses at 23% relative humidity were similar among non-copper surfaces and ranged from 4.5 to 5.9 hours. Assessment of H1N1pdm09 longevity on nonporous surfaces revealed that virus persistence was governed more by differences among HBE culture donors than by surface material. Our findings highlight the potential role of an individual’s respiratory fluid on viral persistence and could help explain heterogeneity in transmission dynamics.

**IMPORTANCE:** Seasonal epidemics and sporadic pandemics of influenza cause a large public health burden. Although influenza viruses disseminate through the environment in respiratory secretions expelled from infected individuals, they can also be transmitted by contaminated surfaces where virus-laden expulsions can deposit. Understanding virus stability on surfaces within the indoor environment is critical to assessing influenza transmission risk. We found that influenza virus stability is affected by the host respiratory secretion in which the virus is expelled, the surface material on which the droplet lands, and the ambient relative humidity of the environment. Influenza viruses can remain infectious on many common surfaces for prolonged periods, with half-lives of 4.5-5.9 hours. These data imply that influenza viruses are highly persistent in indoor environments in biologically-relevant matrices. Decontamination and engineering controls should be used to mitigate influenza virus transmission.

## INTRODUCTION

In 2009, an H1N1 influenza virus emerged from swine and caused a pandemic, with 60.8 million cases in the United States in the first year (*1*). After its emergence, this 2009 H1N1 pandemic virus (H1N1pdm09) became a seasonal influenza virus and it continues to circulate globally (*2*). Seasonal influenza epidemics impose a large public health burden; the Centers for Disease Control and Prevention (CDC) estimated that there were 35 million flu-related illnesses and 20,000 deaths in the United States during the 2019-2020 season (https://www.cdc.gov/flu/about/burden/2019-2020.html). This seasonal burden and the threat of future influenza virus pandemics make it critical to develop strategies to impede influenza virus transmission.

Influenza viruses can spread when infected human hosts expel virus-laden respiratory secretions by breathing, talking, sneezing, or coughing (*3, 4*). Expelled viruses can contaminate surfaces or objects, and these fomites can infect susceptible individuals—a form of indirect contact transmission (*5-7*). Fomites are ubiquitous in household and healthcare settings. Additionally, the demonstrated environmental stability of infectious influenza viruses in laboratory studies make fomite transmission of influenza a contributor to the public health burden of influenza viruses (*8-13*).

Many factors can influence the stability of influenza virus (and therefore, its infectivity) on surfaces, including fomite material. Influenza viruses deposited on porous surfaces, such as fabrics and wood, exhibit reduced persistence of infectivity compared to those on nonporous surfaces, including steel and plastic (*8, 12, 14, 15*). However, influenza virus stability on nonporous surfaces can also depend upon material type; notably, stability on copper is reduced compared to stainless steel (*16*). This observation is consistent with the known antiviral properties of copper (*17, 18*). More studies are required to determine how influenza persists on different materials and could inform engineering controls to mitigate influenza outbreaks.

Environmental conditions such as humidity and temperature also affect the stability of expelled influenza viruses (*10, 11, 19*). Seasonal fluctuations in relative humidity (RH) have been suggested to be a key factor contributing to seasonal patterns of influenza transmission, in part through the impact of RH on influenza virus environmental stability (*20-23*). In addition to changes in virus stability, RH can also influence mucociliary clearance and physico-chemical properties of expelled secretions (*24, 25*). Increased humidity conditions can increase the rate at which virus-laden secretions will fall out of the air and deposit on surfaces. Importantly, respiratory mucus can protect viruses from humidity-mediated decay (*26, 27*), however, it is still unknown whether virus stability in respiratory mucus on different surface materials is influenced by RH.

The matrix composition of droplets is known to influence viral stability on surfaces. Many prior studies examining influenza stability on surfaces have used viral stocks propagated in egg allantoic fluid or tissue culture monolayer cell lines, such as Madin-Darby canine kidney cells. The resultant viral suspensions do not mimic the biochemical composition of virus-laden respiratory secretions expelled from an infected individual. Adding exogenous mucin to cell-grown viral stocks has been reported to have no effect on influenza virus stability (*10, 11*). However, influenza viruses in nasopharyngeal secretions from children with respiratory symptoms were found to be substantially more stable on banknotes than viruses in cell culture medium (*28*). Similarly, our own recent studies have demonstrated that the presence of respiratory mucus from primary human bronchial epithelial (HBE) culture increases the stability of H1N1pdm09 compared to virus suspended in growth media (*26, 27*). These studies indicate that influenza stability needs to be examined further using conditions that more closely mimic respiratory secretions.

While many studies have looked at the individual effects of humidity, surface material, and respiratory droplet composition on influenza infectivity, the interplay between these factors remains understudied. A better understanding of this relationship could improve transmission risk assessment and evidence-based mitigation. In this study, we used H1N1pdm09 viruses grown in HBE cultures from four different patients to better mimic the composition, complexity, and heterogeneity of secretions expelled by infected individuals. HBE cultures are derived from the large airway of explanted human lungs collected during transplant procedures. We then examined the stability of HBE-propagated viruses on nonporous materials under RH conditions ranging from 23% to 98%. We tested six common, nonporous materials: polystyrene (PS) plastic, stainless steel, aluminum, glass, acrylonitrile butadiene styrene (ABS) plastic, and copper. At 23% RH, we found that H1N1pdm09 exhibits similar stability on all the nonporous surfaces tested, except copper. However, at mid-range RH, virus stability is dependent on the surface composition. Additionally, we determined that HBE-propagated H1N1pdm09 is stable over long periods of time on PS plastic, stainless steel, and glass. Importantly, we also found that the HBE patient culture influenced viral stability, suggesting that heterogeneity in human respiratory secretions may contribute to persistence of infectious influenza viruses in the environment.

## RESULTS

### H1N1pdm09 stability in droplets is dependent upon surface material

Environmental factors such as temperature, humidity, and the material of the contaminated surface can impact the survival of influenza viruses (*29, 30*). We propagated H1N1pdm09 in four different HBE patient cultures and collected released virus between 24 and 120 hours post infection (Fig 1A). HBE patient cultures are derived from primary cells collected from explanted human lung tissue, which are differentiated and maintained at an air-liquid interface (*31*). Virus collected from 24-96 hours from each HBE patient culture was pooled and used for subsequent stability analysis. Patient to patient variations in the HBE airway surface liquid could influence the stability of viruses, therefore we performed all studies in all four cultures to look for parameters that influenced virus stability across all cultures.

**Figure 1.**
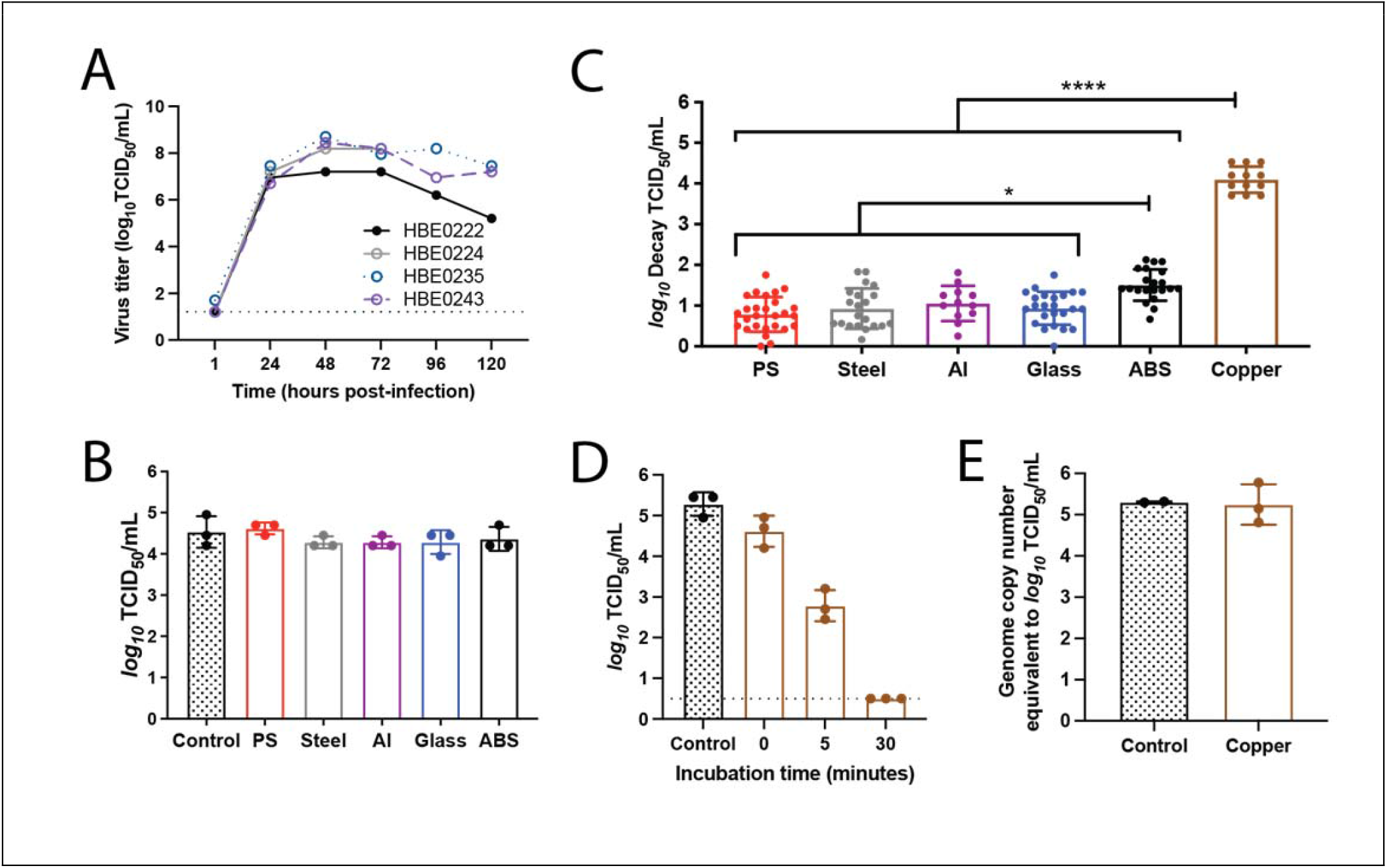
The stability of H1N1pdm09 is dependent on surface material at low relative humidity. **A)** Propagation of H1N1pdm09 in four different HBE patient cultures. Each culture can be related to different pathological states: culture 235 (non-pathological), culture 222 (chronic obstructive pulmonary disease), culture 243 (non-pathological), and culture 224 (idiopathic pulmonary fibrosis). HBE cell cultures were infected with 10^3^ TCID_50_ of virus per well. Virus was collected at the indicated times and titered on MDCK cells. Virus samples from 24 to 96 hours post-infection for each HBE patient culture were then pooled for use in the stability experiments. **B)** Ten 1μL droplets of H1N1pdm09 were deposited on the indicated surfaces and immediately recovered. Virus tittered as in A, each dot represents individual replicates for each HBE culture. **C)** Ten 1μL droplets of H1N1pdm09 (A/CA/07/2009) were incubated at 23% RH on the surface of indicated materials (PS - polystyrene; Al - aluminum; ABS - acrylonitrile butadiene styrene) for two hours. Recovered virus was titered by TCID_50_ assay and viral decay was expressed as the loss of infectivity compared to a control (10μL of virus in a sealed Eppendorf tube outside of the chamber) that was incubated for the same amount of time. Each data point is a replicate of H1N1pdm09 propagated from three to four different HBE cell cultures. Data represent mean values ± standard deviations. Asterisks indicate significantly different mean levels of decay between indicated surface materials, as determined by one-way ANOVA with Tukey’s multiple comparisons test (^*^ p<0.05; ^****^ p<0.0001). No infectious virus was detected after incubation on copper and the *log*_*10*_ decay TCID_50_/mL values were maximal in all replicates. **D)** Ten 1μL droplets of H1N1pdm09 were deposited onto copper. The virus was recovered from the copper surface either immediately after droplet deposition (zero minutes) or after incubation for five or 30 minutes at room RH, which ranged between 41 and 43%. Virus was titered by TCID_50_ assay and data represent mean values ± standard deviations. The dashed line indicates the limit of detection. Data shown is representative of three independent experiments in different HBE cell cultures. **E)** Ten 1μL droplets were recovered following a two-hour incubation on copper at 23% RH. Quantitative PCR for influenza A M gene was used to determine the amount of viral RNA in each sample as compared to a known quantity of viral RNA. Each data point in C-E represents an individual replicate within a study.

To assess virus stability, ten 1 μL droplets of HBE-propagated H1N1pdm09 with a starting titer ranging from 10^6.8^-10^7^ TCID_50_/mL were deposited onto the surface materials of PS plastic, stainless steel, aluminum, glass, ABS plastic, and copper. Droplets were then incubated for two hours inside a desiccator chamber with a RH of 23% to model a dry indoor environment, such as that produced by heating and insulating during a temperate region winter (“flu season”, when influenza spreads efficiently) (*23*). An equivalent volume of virus stock was incubated in a sealed tube for the same amount of time, to serve as a control. Viable virus was quantified by tissue culture infective dose (TCID_50_) assay and compared to the control to determine the amount of decay for each sample. Figure 1B demonstrates the similarity between the control samples and droplets deposited onto each surface and immediately recovered. The overall mean of H1N1pdm09 virus decay on each surface was determined using virus propagated in at least three HBE patient cultures (Fig. 1C).

At 23% RH there was little decay of H1N1pdm09 on PS plastic over the two-hour measurement period (0.79 ± 0.42 *log*_*10*_ TCID_50_/mL) relative to the sealed control, as previously observed (*26*). Similar to PS plastic, the mean values of decay on stainless steel, aluminum, and glass were less than or at 1 *log*_*10*_ TCID_50_/mL. Decay was significantly higher on ABS plastic (1.5 ± 0.39 *log*_*10*_ TCID_50_/mL) and copper, where the deposited virus decayed to undetectable levels (Fig. 1C). The differences in decay did not arise from variable recovery of H1N1pdm09 from the different materials since immediate recovery of droplets from each surface resulted in similar titers to the control (Fig. 1B).

In stark contrast to the moderate decay on ABS plastic after two hours, no viable H1N1pdm09 was detected after deposition on copper at 23% RH in any replicate, where each replicate reached the maximal decay titer (Fig. 1C). To address whether reduced recovery of H1N1pdm09 from copper impacted the magnitude of decay, droplets were spotted onto the surface and then immediately recovered. Immediate recovery of virus from copper revealed similar viral titers to the control sample (Fig. 1D). Additionally, following a two-hour incubation on copper, viral RNA was extracted from recovered virus solution and quantified by qPCR. No difference in total RNA was detected (Fig. 1E), indicating that the rapid inactivation on copper was not due to poor recovery from the material. To determine the longevity of H1N1pdm09 on copper, we shortened the incubation time to five or 30 minutes. Given the short time scale and initial chamber equilibration, H1N1pdm09 droplets were incubated at room RH, which was stable between 41% and 43%. After five minutes on copper, the viability of H1N1pdm09 decreased to 2.78 ± 0.38 *log*_*10*_ TCID_50_/mL, whereas a 30-minute incubation resulted in a titer that was below the limit of detection (Fig. 1D). Taken together, these data indicate that at 23% RH, H1N1pdm09 is most stable on PS plastic, stainless steel, aluminum, and glass, while it is slightly less stable on ABS plastic, and is rapidly inactivated on copper.

### H1N1pdm09 stability is impacted by relative humidity in a surface dependent manner

Prior studies have revealed a relationship between RH and virus stability of enveloped viruses. In many cases, inactivation has been shown to be fastest at mid-range RH (about 40-70%) compared to low (23%) or high (>80%) RH (*19*). However, our previously published data demonstrated that H1N1pdm09 is protected from RH-mediated decay in droplets supplemented with airway surface liquid from HBE patient cultures on PS plastic (*26*). To determine how the stability of HBE-propagated H1N1pdm09 on different surface materials is affected by RH, we spotted 1μL droplets of H1N1pdm09 on each of the six nonporous surfaces over a range of RH conditions and incubated the droplets for two hours. In addition to 23% RH shown in Figure 1, we tested two mid-range RHs (43% and 55%), which are typical of indoor environments during summer in temperate regions, and one high RH (98%), which mimics conditions during rainy periods and in airways. Viable H1N1pdm09 was recovered from PS plastic, stainless steel, aluminum, glass, and ABS plastic at all RHs tested, but the magnitude of decay varied by both RH and surface material (Fig. 2A-E). Overall, little decay was observed on PS plastic and glass at 23%, 43%, and 55% RH, with 98% RH being more stable than 43% and 55% RH (Fig. 2A and 2D). H1N1pdm09 displayed maximal decay on stainless steel and aluminum at the mid-range RHs, but was more stable at both 23% and 98% RH (Fig. 2B-C). Decay of H1N1pdm09 on ABS plastic was the highest at mid-range humidities, followed by 23% RH, which showed significantly more decay than 98% RH (Fig. 2E, Table 1). Similar to what was seen at 23% RH, no infectious virus was detected on copper surfaces after two hours under all other RH conditions (Fig 2F).

**Table 1.**
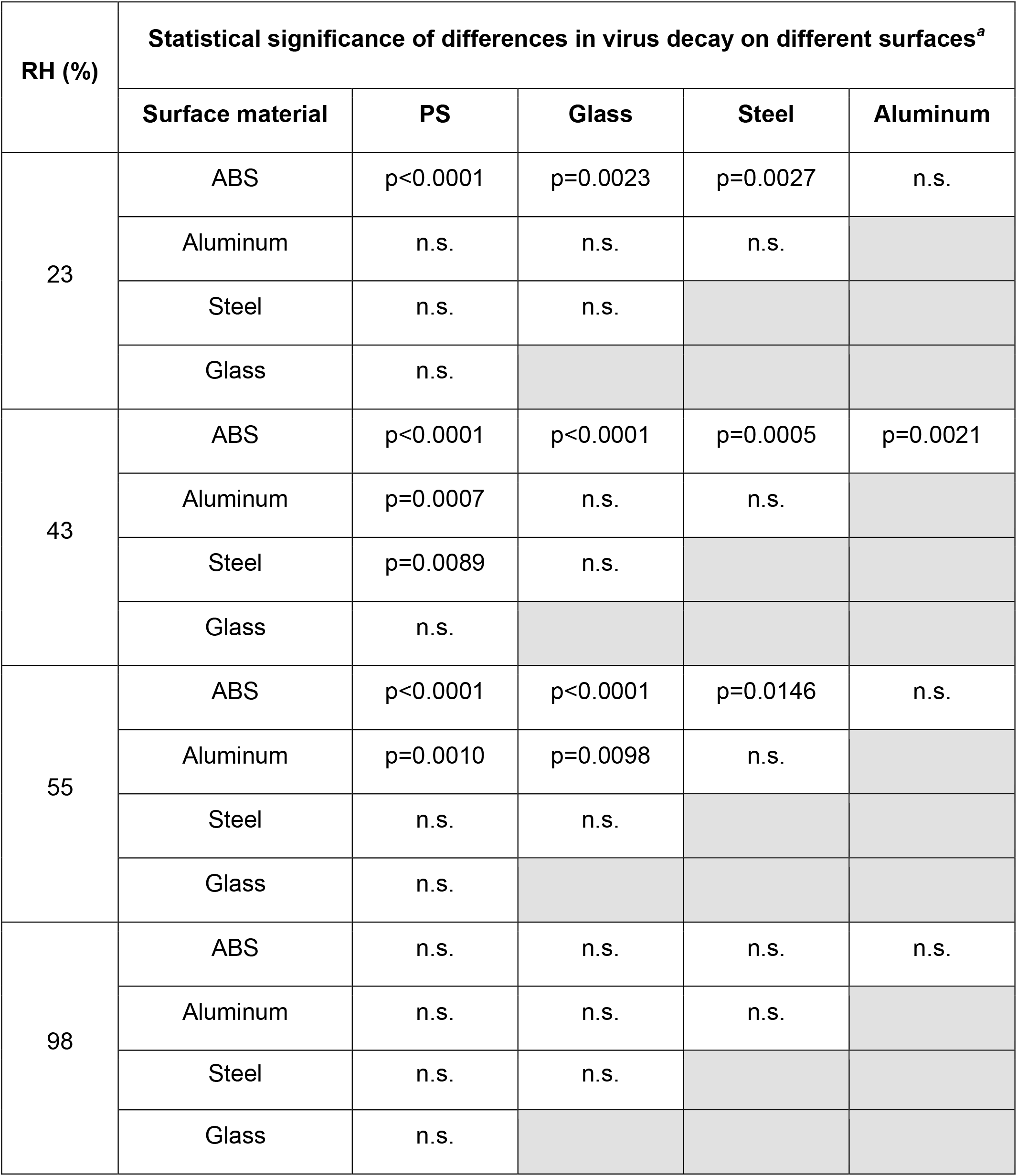
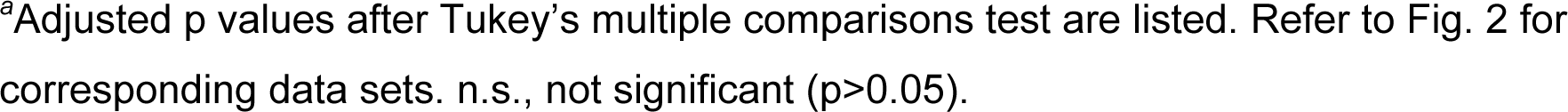
Statistical analysis of the surface-based variations in the stability of H1N1pdm09 in droplets at different RH.

| RH (%) | Statistical significance of differences in virus decay on different surfaces<sup>a</sup> |  |  |  |  |
| --- | --- | --- | --- | --- | --- |
|  | Surface material | PS | Glass | Steel | Aluminum |
| 23 | ABS | p<0.0001 | p=0.0023 | p=0.0027 | n.s. |
|  | Aluminum | n.s. | n.s. | n.s. |  |
|  | Steel | n.s. | n.s. |  |  |
|  | Glass | n.s. |  |  |  |
| 43 | ABS | p<0.0001 | p<0.0001 | p=0.0005 | p=0.0021 |
|  | Aluminum | p=0.0007 | n.s. | n.s. |  |
|  | Steel | p=0.0089 | n.s. |  |  |
|  | Glass | n.s. |  |  |  |
| 55 | ABS | p<0.0001 | p<0.0001 | p=0.0146 | n.s. |
|  | Aluminum | p=0.0010 | p=0.0098 | n.s. |  |
|  | Steel | n.s. | n.s. |  |  |
|  | Glass | n.s. |  |  |  |
| 98 | ABS | n.s. | n.s. | n.s. | n.s. |
|  | Aluminum | n.s. | n.s. | n.s. |  |
|  | Steel | n.s. | n.s. |  |  |
|  | Glass | n.s. |  |  |  |

<sup>a</sup>Adjusted p values after Tukey's multiple comparisons test are listed. Refer to Fig. 2 for corresponding data sets. n.s., not significant (p>0.05).

**Figure 2.**
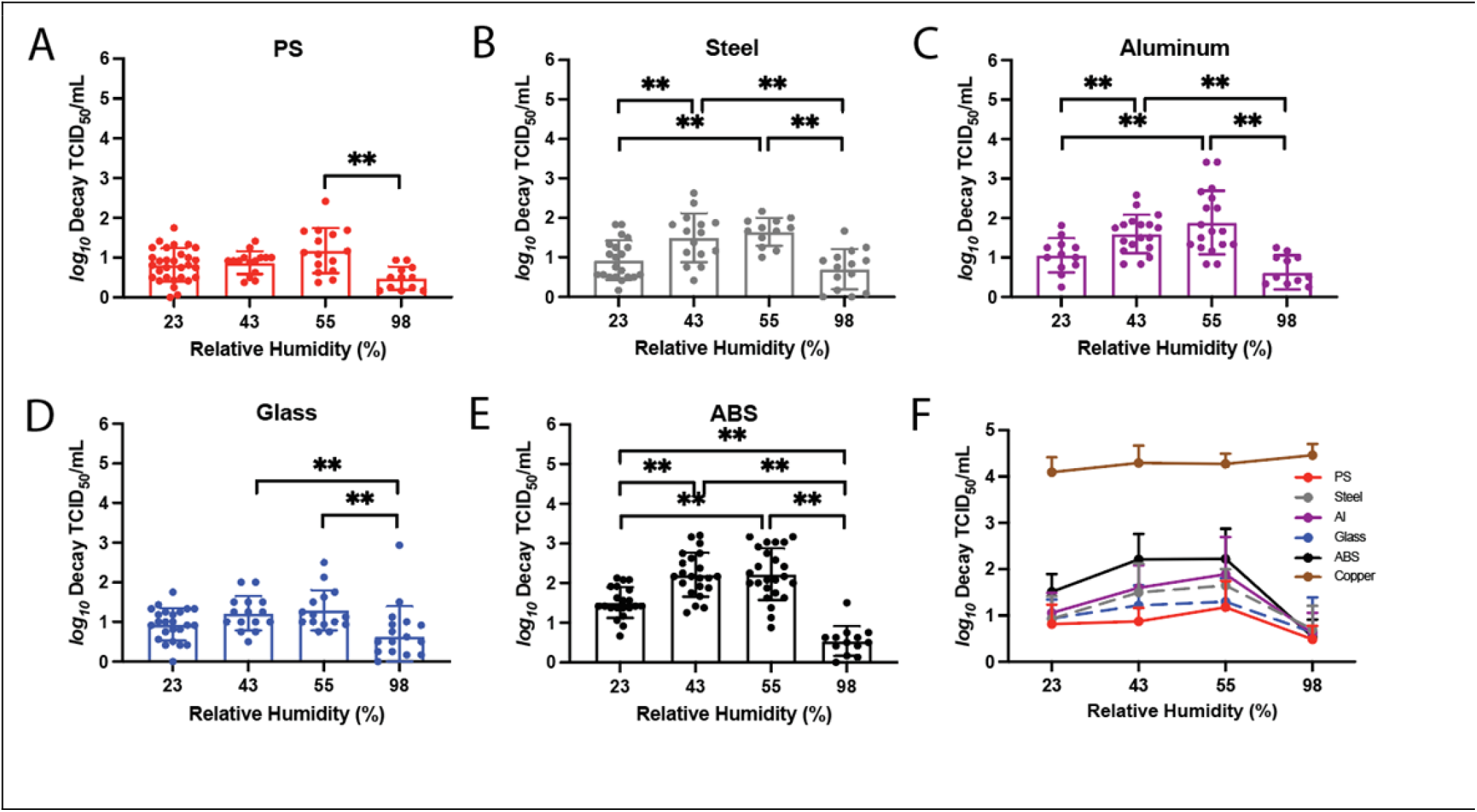
Impact of RH on H1N1pdm09 stability is surface dependent. The stability of H1N1pdm09 in 10 1μL droplets on each material was tested under a range of RH conditions: 23%, 43%, 55%, and 98%. The infectivity decay of the virus after two hours on **A)** PS plastic, **B)** stainless steel, **C)** aluminum, **D)** glass, and **E)** ABS plastic was calculated as previously described in figure 1. Virus propagated from at least three different HBE cultures was tested under each condition. Each data point represents a single replicate, and the results are shown with means and standard deviations. Two-way ANOVA, using surface material and RH as the variables, with Tukey’s multiple comparisons test was performed on the results in **A-E** to analyze the effect of RH on the virus decay: ^**^, p<0.01. **F)** The mean ± standard deviation of virus decay on each of the surfaces in A to E and copper were plotted against the relative humidity (RH).

Direct comparison of virus decay by surface reveals a hierarchy of viral persistence by surface type (Fig. 2F). Viruses appear most stable on PS plastic and glass surfaces, with increased decay on stainless steel and aluminum, followed by ABS plastic. No surface-based differences in virus decay were observed at 98% RH where the mean decay values were all below 1 log10TCID50/mL, even on ABS plastic (Fig. 2F, Table 1). Taken together, our results demonstrate that RH can affect the stability of HBE-propagated H1N1pdm09, but this feature was only apparent on certain surface materials. Additionally, certain nonporous materials exhibit more viral destabilization than others.

### Predicted half-life of virus depends more strongly on HBE culture than on surface material

Our experiments used virus propagated from different HBE cultures derived from individual patients (Fig 1A). Patient-to-patient variability in the HBE airway surface liquid surrounding the released virions might also contribute to differences in viral stability on various materials. Therefore, we wanted to examine the relationship between surface material and HBE composition on virus stability. To do this, we spotted HBE-propagated H1N1pdm09 droplets (from the four distinct patient cultures) for two, eight, or 24 hours on PS plastic, stainless steel, glass, and ABS plastic. To mimic a dry indoor environment in winter, H1N1pdm09 droplets were incubated in a chamber at 23% RH. The recorded RH inside the chamber was stable for up to 24 hours after a brief equilibration period during the first few minutes. Viral titers were assessed after recovery of the droplets at each time interval, and then used to calculate the half-life of each HBE-grown virus.

To estimate the half-life of viable influenza virus and its dependence on surface and HBE culture, we used a hierarchical regression model adapted from our own previously published Bayesian statistical methods for analyzing viral environmental stability (*32, 33*). Briefly, we inferred virus half-lives and surface- and culture-level effects upon those half-lives directly from raw titration data (inoculated wells positive or negative for virus infection). Initially, we analyzed all the data together, using a model that assumes that surface and culture act independently (and multiplicatively) to modify the half-life. The models quantify virus from positive or negative readouts of inoculated wells by treating well inoculation as a “single-hit” process, with an assumed Poisson distribution for the number of virions. This is similar to traditional endpoint titration statistical approaches such as the Spearman-Karber and Reed-Muensch methods. However, implementing this in a Bayesian framework connects our model directly to a regression for estimating half-lives, and we can obtain principled estimates of uncertainty (such as 95% credible intervals) for each individual titration, rather than just a single number. The resulting model fits for each culture and surface are displayed in figure 3, and the resultant estimates of culture and surface effects in figure 4.

**Figure 3.**
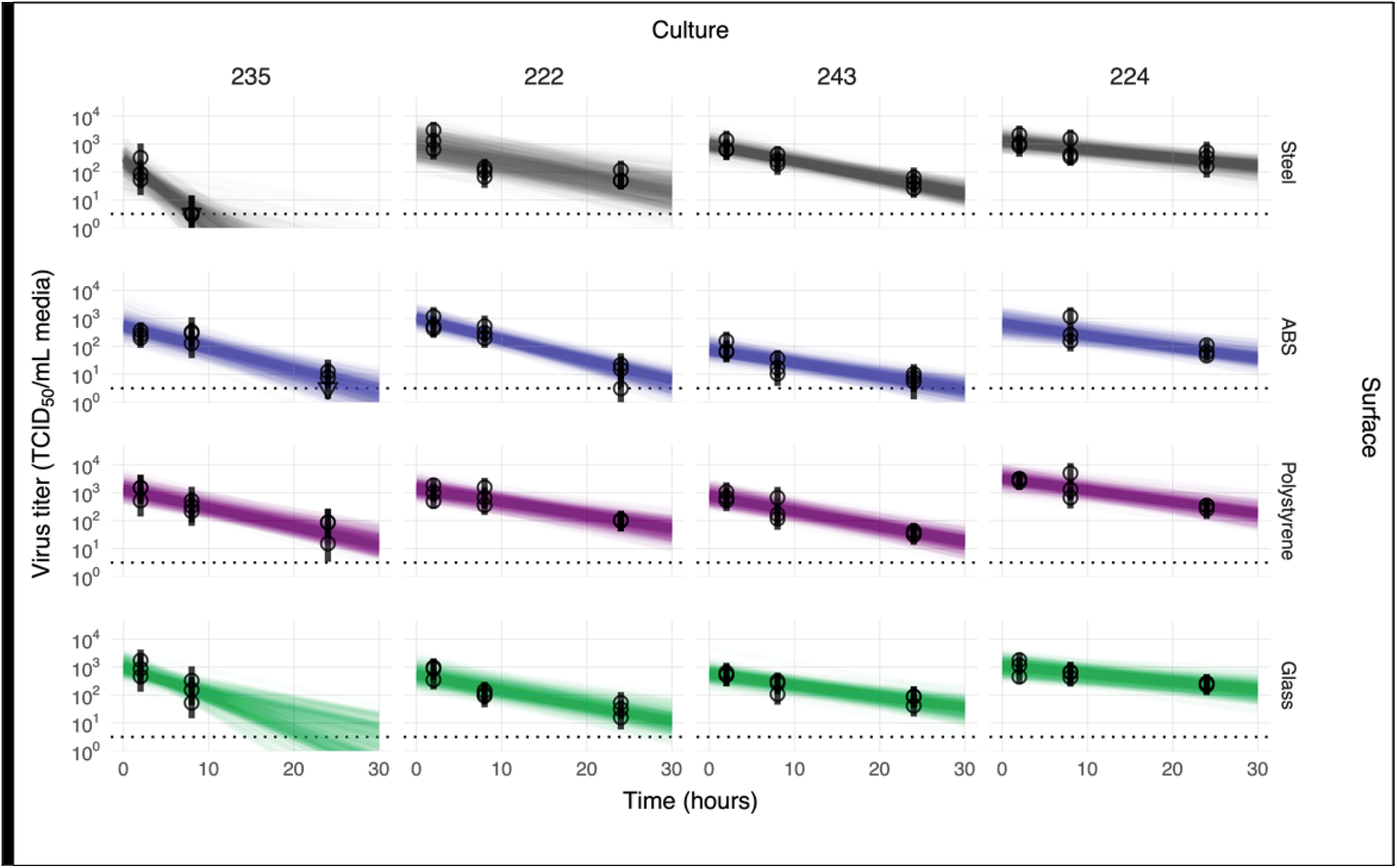
Linear regression of viral stability on surfaces at 23% RH over time by HBE culture to estimate surface and culture effects. Regression lines representing predicted exponential decay of log_10_ virus titer over time are plotted alongside measured (directly-inferred) virus titers. Predicted decay reflects the estimated effects of surface (row) and culture (column). For each experiment (surface / HBE culture pair), semitransparent regression lines visualize the inferred joint posterior distribution of the virus exponential decay rate and the individual sample intercepts (i.e. virus titers at t = 0, which can vary about the mean initial titer for the experiment). 50 random posterior draws are shown. To visualize the estimated variation in initial titers (intercepts), 6 lines are plotted per experiment per draw, one for each of 6 randomly-chosen titers from that experiment (since each titer has its own estimated t = 0 value). This yields 300 plotted lines per experiment. A new set of 6 random titers per experiment is chosen for each draw. Points with black bars show individually estimated titer values (point: posterior median titer estimate; bar: 95% credible interval). Samples with no positive titration wells are plotted as triangles at the approximate LOD (dotted horizontal line).

**Figure 4.**
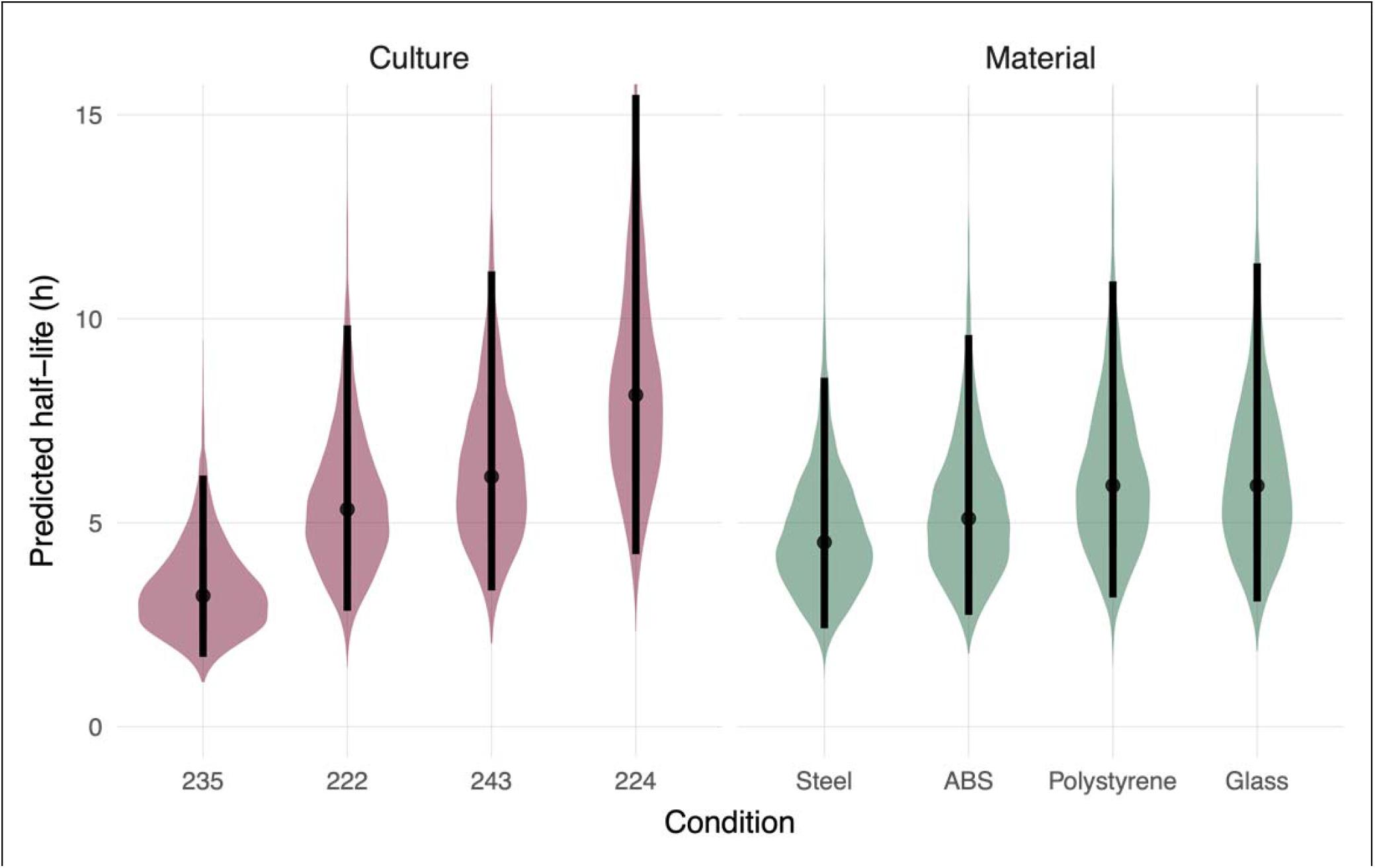
Predicted half-life based on surface material or patient-derived culture. Violin plots indicate the posterior distributions of the half-life of viable virus based on the estimated exponential decay rates of the virus in different HBE cultures on a “neutral” surface (left side) and on surfaces with a “neutral” HBE culture (right side), thus separating out culture and surface effects. Dots indicate the posterior median estimates and the black lines indicate a 95% credible interval (CI). Values for the median half-lives and 95% CI are presented in Table 2.

To estimate virus half-lives, we coupled our titer estimation model to a simple regression model in which virus is assumed to decay exponentially over time. The slope of individual regression lines predicting titer as a function of time will provide an estimate of the exponential decay rate, which can be converted directly into a half-life. To estimate the effects of individual surfaces and cultures on virus persistence, we used a regression model in which we treated log half-life of the virus in a given culture on a given surface as depending on three quantities: a “culture effect” (assumed to apply across surfaces), a “surface effect” (assumed to apply across cultures), and a single intercept (representing the average log half-life across all cultures and surfaces tested). This simple model assumes that the culture effect and surface effect act independently to modify the half-life of the virus.

**Table 2:** Virus half-life by surface material and HBE culture

| Virus Half-Life by Surface Material<br>median [95% credible interval] |  |  |  | Virus Half-Life by HBE Culture<br>median [95% credible interval] |  |  |  |
| --- | --- | --- | --- | --- | --- | --- | --- |
| Stainless<br>Steel | ABS<br>Plastic | PS<br>Plastic | Glass | HBE 235 | HBE 222 | HBE 243 | HBE 224 |
| 4.52<br>[2.41, 8.56] | 5.10<br>[2.74, 9.60] | 5.91<br>[3.17, 10.9] | 5.91<br>[3.07, 11.4] | 3.21<br>[1.71, 6.16] | 5.33<br>[2.84, 9.84] | 6.13<br>[3.34, 11.2] | 8.13<br>[4.23, 15.5] |

Using this approach on our experimental data, we report the predicted half-life for the virus in each given culture on a hypothetical “neutral” (average) surface—that is, with zero surface effect. Similarly, we report estimated surface effects as predicted half-lives on each surface given a hypothetical “neutral” (average) culture—that is, with zero culture effect (Fig 4 and Table 2). The median viral half-life of a neutral culture on different surface materials (excluding copper) ranged between 4.5 and 5.9 hours (Table 2). In contrast, the median viral half-life range on a neutral surface broken down by HBE culture was 3.2 and 8.1 hours (Table 2). The greater estimated variation in half-lives by HBE culture than by surface material, suggests that host-specific variation may be an important determinant of environmental transmission risk. Calculations of virus half-life for each HBE culture include data from all surface materials (excluding copper). However, given the similarity of virus half-life values across surface material type, it is likely that the large variations observed between virus half-life by HBE culture are not due to surface material but rather differences in the airway secretion composition.

## DISCUSSION

In this study, we investigated influenza virus stability on different types of surfaces using H1N1pdm09 propagated from three-dimensional patient-derived lung cultures. Stability of H1N1pdm09 in the presence of airway surface liquid, appears to be surface-dependent under various common indoor RH conditions tested (23%, 43% and 55%). The notable exception was deposition on copper, which completely inactivated virus at all RH conditions and as quickly as 30 minutes at room RH. Of the other nonporous surfaces tested (excluding copper), the half-life of HBE-propagated H1N1pdm09 at 23% was similar on PS plastic, stainless steel, aluminum, and glass. Interestingly, a subsequent analysis revealed that the half-life of infectious H1N1pdm09 at 23% RH varied substantially among the different HBE patient cultures that were used to propagate the virus. This final observation suggests that the respiratory secretion composition may have a profound impact on the persistence of viruses in the environment, and that individual variation may matter a great deal.

The rapid inactivation of HBE-propagated H1N1pdm09 observed on copper at all RH conditions tested indicates that this effect is mediated by the surface itself and is robust against other environmental conditions. Noyce et al. have also described inactivation of 20 μL droplets of A/PR/8/34 (H1N1) after six hours on a copper surface (*16*), whereas in our study, 1μL droplets of H1N1pdm09 had no detectable infectious virus after just 30 minutes. The size and composition of the influenza virus-containing droplets deposited on a given surface may impact virus viability and decay kinetics. Virucidal activity of copper and copper alloys has been reported for seasonal coronaviruses, SARS-CoV-2, and norovirus (*17, 34-37*). The potent antiviral properties of copper have been shown by Warnes *et al* to damage membrane and surface proteins, and to produce nonspecific fragmentation of the coronavirus 229E genome (*18*). Previous studies have also shown that bacteria are rapidly killed on dry copper surfaces (*38*), which has been proposed to be mediated by DNA damage (*39, 40*). Addition of copper oxide or copper nanocompounds to frequently touched surfaces could reduce the spread of influenza (*41-43*).

Environmental conditions such as RH could affect the transmissibility of influenza virus fomites by influencing virus stability. Relevant RH conditions include those similar to indoor RH during temperate zone winters (approximately 20%), and those similar to indoor RH during temperate zone summers (40-60%). Previous work has suggested a U-shaped relationship between RH and virus stability for enveloped viruses including SARS-CoV-2 and influenza, where virus stability is lowest at mid-range RH conditions (*19, 33*). However, our studies with influenza viruses suspended in HBE airway surface liquid have revealed that viruses are protected from RH-mediated decay on PS plastic (*26*). In this study, we reproduced our previous observations that HBE-propagated H1N1pdm09 was stable on PS plastic (Fig 2). However, we also observed variable stability of H1N1pdm09 propagated in HBE patient cultures at 23%, 43%, and 55% RH on stainless steel, aluminum, and ABS plastic nonporous surfaces (Fig 2). These data suggest that RH-mediated decay of influenza may be more pronounced on different surfaces and may also depend on the composition of respiratory droplet.

Most surprisingly, HBE-propagated influenza virus half-life at 23% RH appeared similar across surfaces, but droplet composition from HBE cultures impacted H1N1pdm09 stability more (Fig 4). This suggests a key role for droplet composition on the persistence of viruses in the environment. Our cultures included those from normal (non-pathological) lung transplants (culture 235 and 243) and those with disease states (222 and 224). More experiments utilizing virus grown from greater numbers of HBE cultures will be required to determine if and how pathological states contribute to virus stability. Additionally, further studies examining the differences in expelled droplet composition and volume across individuals could yield insight into the observed heterogeneity of influenza transmission (*44*). The SARS-CoV-2 pandemic was driven by cluster-based transmission: 20% of infected individuals seeded 80% of secondary infections (*45*). This suggests that individual characteristics of ‘superspreaders’ may impact the emergence of novel respiratory viruses (*46, 47*). Understanding which respiratory secretion components are beneficial or harmful to virus stability, and how these map onto identifiable traits of human individuals, could aid in understanding what makes a ‘superspreader’.

The emergence of multiple respiratory pandemic viruses in the last few decades has highlighted the importance of understanding virus transmission pathways (inhalation of aerosols, spray of large droplets, touching of fomites), and determining the impact of virus traits, environmental conditions, and host factors on each mode (*48*). Persistence of infectious influenza virus on various non-porous surfaces over extended time periods indicates that regular decontamination of frequently-touched surfaces, in combination with engineering controls such as indoor RH control, could be effective nonpharmaceutical interventions to limit fomite transmission of influenza viruses. Patterns of heterogeneous virus stability by individual patient culture suggest a potential contributing mechanism to observed host heterogeneity in virus transmission.

## MATERIALS AND METHODS

### Cells and virus

Madin-Darby canine kidney (MDCK) cells (ATCC) were maintained in Eagle’s Minimum Essential Medium supplemented with 10% fetal bovine serum, 2 mM L-glutamine, and 1% penicillin/streptomycin. Four primary human bronchial epithelial cell (HBE) cultures were differentiated from lung tissues of four different patients and were maintained at an air-liquid interface in transwells (*31*). Each HBE culture was derived from patient lungs with various pathological states: culture 235 (non-pathological), culture 222 (chronic obstructive pulmonary disease), culture 243 (non-pathological), and culture 224 (idiopathic pulmonary fibrosis).

The 2009 pandemic H1N1 influenza A virus, A/California/07/2009 (H1N1pdm09), was acquired as previously described (*26*). To prepare HBE-propagated virus stock, 3 *log*_*10*_ TCID_50_ of virus was added to each transwell from the apical side after the cells were washed once with phosphate buffered saline (PBS). The inoculum was removed after 1 hour and the HBE-propagated virus was collected with 150mL of PBS every 24 hours up to five times. Viral washes collected at the same time from different transwells of the same HBE cell culture were pooled together and titered on MDCK cells. Up to four viral washes with the highest titers from the same HBE culture were combined. To make an abundant HBE-propagated virus stock without diluting physiological components in the suspension, the combined virus wash was 1:1 diluted with the cell wash from the same HBE culture, which had been collected prior to infection. To account for patient specific variations, four stocks of HBE-propagated virus from different cultures were made. The stocks were aliquoted and stored at -80 °C before use.

### Surface preparation

Six surface materials were selected because of common use: polystyrene (PS) plastic, stainless steel, aluminum, glass, acrylonitrile butadiene styrene (ABS) plastic, and copper. Disposable PS plastic was readily available as the flat bottom of a 6-well tissue culture plate (TPP^®^, Sigma). Disposable glass cover slides (22CIR-1, Thermo Fisher) were used as glass surfaces without additional preparation. Circular and smooth surface coupons of 304 stainless steel (20 GA), 6061 aluminum (1.6 mm thick), ABS plastic (black, 0.6 mm thick), and 110 copper (0.6 mm thick) were purchased through Alumagraphics with a uniform diameter of 20 mm. The coupons were cleaned with an interfering-residual-free detergent (Alconox^®^) prior to sterilization (stainless steel, aluminum, glass) or disinfection with isopropanol (ABS plastic). The coupons were reused after cleaning and sterilization (or disinfection) following the assay.

### Virus stability assay

Saturated saline solutions were placed in a sealed chamber to condition the interior relative humidity (RH) to desired levels – KCH_3_COO for 23%, K_2_CO_3_ for 43%, Mg(NO_3_)_2_ for 55%, and K_2_SO_4_ for 98% (*19*). The temperature and RH inside the chamber were monitored by a HOBO^®^ logger. Ten 1μL droplets of the HBE-propagated virus were pipetted either directly onto each well (PS plastic) or onto a coupon inside a lidless 6-well plate in triplicates. The plate was immediately transferred into the RH chamber upon droplet deposition. After incubation for 0.5, two, eight, or 24 hour(s), the virus was collected by rinsing each well, or each coupon, with 1 mL Leibovitz’s L-15 Medium, resulting in a 1:100 dilution compared with the volume of the deposited droplets. The control sample was a bulk 10μL suspension of the same HBE-propagated virus in a 1.5-mL Eppendorf tube, which was incubated for the same amount of time without RH chamber conditioning or contact with the coupons. The RH chamber was not used for shorter virus incubation on copper coupons since the chamber required at least five minutes to equilibrate the interior RH. Instead, the virus was incubated on copper coupons inside a lidless 6-well plate at room RH. Either immediately (zero minutes) or after five or 30 minutes of incubation, the virus was recovered as previously described. The control sample was immediately diluted without any incubation. All procedures were conducted inside a biosafety cabinet at room temperature (22-24°C). Under each condition, HBE-propagated influenza H1N1pdm09 from at least three different cultures was tested.

The decay of H1N1pdm09 estimated in Figs. 1 and 2 is defined as loss of infectivity as determined by TCID_50_ assay (*49*). Specifically, the virus samples were titered by 10-fold serial dilutions on a 96-well plate of confluent MDCK cells in quadruplicates. A 24-well plate was also used where undiluted virus samples were incubated on MDCK cells in duplicates to provide the limit of detection at 0.5 *log*_*10*_ TCID_50_/mL. The loss of virus infectivity was then calculated by comparing titers of the virus on the surface coupons with titers of the control samples incubated for the same amount of time:

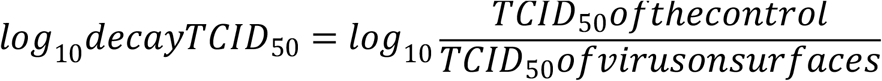

The results of the virus decay were analyzed on GraphPad Prism 8. Statistical analysis excluded results from copper because the virus decay was mostly beyond the detection limit. At 23% RH, one-way analysis of variance (ANOVA) using surface material as the variable was performed with Tukey’s multiple comparisons test to define surface-based variations in virus decay within two hours. Two-way ANOVA using RH (or incubation time) as the second variable was performed with Tukey’s multiple comparisons test to define the effect of RH (or incubation time) on virus decay. Due to the interactive effects of RH and surface material on virus decay, the analysis of surface-based variations at different RH was adjusted with Tukey’s multiple comparison tests following two-way ANOVA and summarized in Table 1.

### Quantitative reverse transcription polymerase chain reaction (qRT-PCR)

The RNA of H1N1pdm09 was extracted from virus samples using a PureLink^™^ viral RNA/DNA mini kit (Invitrogen). For qRT-PCR, 5μL of the viral RNA was analyzed using a TaqMan^™^ RNA-to-C_T_^™^ 1-step kit (Thermo Fisher). Primers specific to the matrix gene (M) were used at a concentration of 0.1 μM—M25 F (5’-AGA TGA GTC TTC TAA CCG AGG TCG-3’) and M123 R (5’-GC AAA GAC ATC TTC AAG TCT CTG-3’)—and an M64 probe (FAM-TCA GGC CCC CTC AAA GCC GA-NFQ) at 0.25 μM. All reactions were performed in duplicate. The thermal cycling step was conducted on a StepOnePlus^™^ Real-Time PCR System (Applied Biosystems) with StepOne^™^ Software (Version 2.3) following manufacturer’s recommendations. A standard curve was generated with five 10-fold serial dilutions of viral RNA extracted from a stock virus with a known TCID_50_ titer.

### Bayesian inference methods

To estimate the effects of surface and culture on virus half-lives (Figs. 3, 4), we jointly inferred these effects and the corresponding half-lives directly from raw titration data (inoculated wells positive or negative for virus infection), using Bayesian inference with suite of custom statistical models. Below, we provide a conceptual overview of those models, followed by a full technical description. We have also published code implementing the model for reproducibility.

The models quantify virus from positive or negative readouts of inoculated wells by treating well inoculation as a “single-hit” process: if at least one virion successfully infects a cell within the well, the well will show evidence of infection. We then estimate the virus concentration in TCID_50_ from the distribution of positive and negative wells observed at various dilutions of the sample. We assume that the number of virions inoculated into any given well is Poisson-distributed, with a mean given by the virus concentration in the diluted sample used for that well. This is the same “Poisson single-hit” assumption that underlies traditional endpoint titration statistical approaches such as the Spearman-Karber and Reed-Muensch methods. The difference is that by implementing this in a Bayesian framework, we are able to connect our model directly to a regression for estimating half-lives, and we are able to obtain principled estimates of uncertainty (such as 95% credible intervals) for each estimated quantity, rather than just a single number.

To estimate virus half-lives, we couple our titer estimation model to a simple regression model in which virus is assumed to decay exponentially over time. The slope of a regression line predicting log titer as a function of time therefore gives an estimate of the exponential decay rate, which can be converted directly into a half-life. Our Bayesian approach allows us to account in a principled way for various sources of noise, such as variation in the initial quantities of virus deposited onto individual surface coupons.

Finally, to estimate the effects of individual surfaces and cultures on virus persistence, we used a regression model in which we treated log half-life of the virus in a given culture on a given surface as depending on three quantities: a “culture effect” (assumed to apply across surfaces), a “surface effect” (assumed to apply across cultures), and a single intercept (representing the average log half-life across all cultures and surfaces tested). The model assumes that the culture effect and surface effect act independently to modify the half-life of the virus. We estimated these culture and surface effects, and the intercept, from our data.

To aid interpretability, we report estimated culture effects in units of half-life. Specifically, we report the predicted half-life for the virus in the given culture on a “neutral” (average) surface—that is, with zero surface effect. Similarly, we report estimated surface effects as predicted half-lives on the given surface given a “neutral” (average) culture—that is, with zero culture effect.

#### Notation

In the model notation that follows, the symbol ∼ denotes that a random variable is distributed according to the given distribution. Normal distributions are parametrized as:

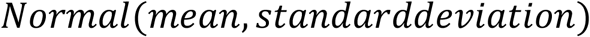

Positive-constrained Normal distributions (“Pos-Normal”) are parametrized as:

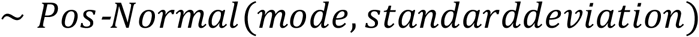

Beta distributions are parameterized as

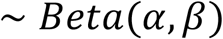

with canonical shape parameters *α, β* > 0 (which can be thought of as 1 plus the number of prior successes and failures, respectively, in a set of *α* + *β* binomial trials)

#### Titer inference and Model description

We inferred individual titers directly from titration well data according to a Poisson single-hit model (*32*), as described in (*33*).

Briefly, we assume that individual wells for a sample *i* are positive if at least one virion successfully infects a cell. We assume the number of virions that successfully infect cells within a given well is Poisson distributed with a mean given by the concentration of viable virions in the plated sample.

This gives us our likelihood function, assuming independence of outcomes across wells. Titrated doses introduced to each cell-culture well were of volume 0.1 mL, so we incremented inferred titers by 1 to convert to units of *log*_*10*_*TCID*_50_/[mL].

The one variation from the model described in (*33*) is that here we also had negative control wells. We used these to estimate the probability of a false positive well. Whereas in (*33*) the mean of our Poisson was given by

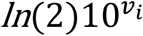

where *v*_*i*_ is the concentration of viable virus in *TCID*_50_/[0.1 mL], here we have a mean of

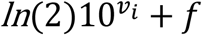

where *f* is a constant governing the false positive rate and is related to the probability of a false positive by:

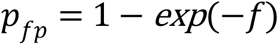

so:

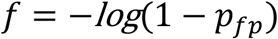

#### Prior distributions

We assigned a weakly informative Normal prior to the *log10* titers *v*_*i*_ (*v*_*i*_ is the titer for sample *i* measured in *log*_*10*_*TCID*_50_/[0.1 mL], since wells were inoculated with 0.1*mL*).

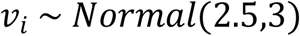

We placed a Beta prior on the false positive probability *p*_*fp*_, assuming it to be small but allowing it to be non-trivial:

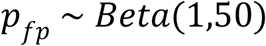

#### Predictive checks

We assessed the appropriateness of prior distribution choices using prior predictive checks. The prior checks suggested that prior distributions were agnostic over the titer values of interest.

### Half-life inference model

#### Model description

To infer half-lives of viable virus in the various experiments, we used a regression model.

For each experimental condition *i*, we have two sets of measurements: treatment samples deposited on surfaces and incubated at a given temperature and humidity, and control samples kept in bulk solution at room temperature. We assume that each sample *j* for experimental condition *i*, whether treatment or control, had some unknown initial *log*_*10*_ titer value *v*_*ij*0_ at *t* = 0.

We assume that all of these initial values are Normally distributed about a mean initial *log* titer 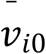 for the experiment, with an unknown experiment-specific standard deviation *σ*_*vi*_:

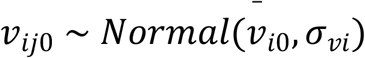

We modeled loss of viable virus as exponential decay at a rate *δ*_*i*_ for treatment samples in condition *i* and *λ*_*i*_ for the corresponding control samples. For the treatment samples, we assumed there was an experiment-specific mean amount of virus lost during deposition *ℓ*_*i*_.

It follows that the quantity *v*_*ij*_ of virus sampled at time *t*_*ij*_ is given by

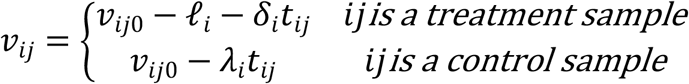

We then used the direct-from-well data likelihood function described above, except that instead of estimating individual titers independently, we estimated the values of *δ*_*i*_, *λ*_*i*_, and *ℓ*_*i*_ under the assumption that our observed well data reflected the corresponding predicted titers *v*_*ij*_.

#### Prior distributions

We placed a weakly informative Normal prior on the mean initial *log*_*10*_ titers 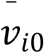 to reflect the known inocula:

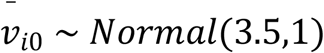

We placed a Pos-Normal prior on the initial titer standard deviations *σ*_*vi*_:

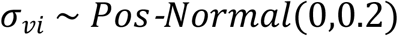

This allows either for large variation (more than *±* 0.5 *log*_*10*_) about the experiment mean or for substantially less variation, depending on the data.

We placed Normal priors on the log treatment and control half-lives *Δ*_*i*_ and *Λ*_*i*_, where 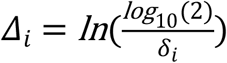 and 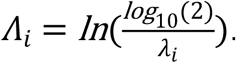. We made the priors weakly informative (diffuse over the biologically plausible half-lives); we verified this with prior predictive checks.

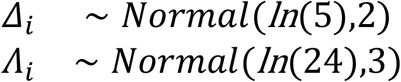

We placed a Pos-Normal prior on the experiment mean deposition losses *ℓ*_*i*_:

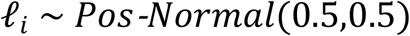

The prior for the false positive probability *p*_*fp*_ was as in the titer estimation model (see ‘Titer inference and Model description’).

#### Predictive checks

We assessed the appropriateness of prior distribution choices using prior predictive checks and assessed goodness of fit for the estimated model using posterior predictive checks.

### Model to estimate culture and surface effects

#### Model description

To estimate and compare the effects of surface and culture on virus persistence, we used a simple regression model. We estimated treatment and control log half-lives *Δ*_*i*_ and *Λ*_*i*_ as in the half-life inference model (detailed above), but instead of placing priors directly on them, we assumed that the log half-lives *Δ*_*i*_ could be predicted according to a linear equation with intercept 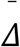, culture-specific effects *X*_*n*_, and surface-specific effects *μ*_*m*_:

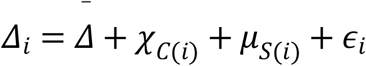

where 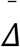 is an intercept representing the log half-life for virus in a neutral (0-effect) culture on a neutral (0-effect) surface, *C*(*i*) is the culture for experiment *i, S*(*i*) is the surface for experiment *i*, and *ε*_*i*_ is a Normally distributed error term with an estimated standard deviation *σ*_*Δ*_:

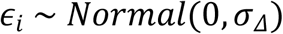

We assumed that the control log half-lives *Λ*_*i*_ were Normally distributed about an unknown mean *Λ* with an unknown standard deviation *σ*_*Λ*_:

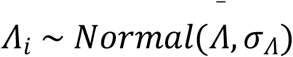

#### Prior distributions

We placed Normal priors on the treatment intercept log half-life 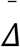 and the mean control half-life 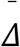 equivalent to the normal priors for *Δ*_*i*_ and *Λ*_*i*_ in the half-life estimation model

(eqn. <u>[eqn:log-half-life-priors]</u>, sec. 3).

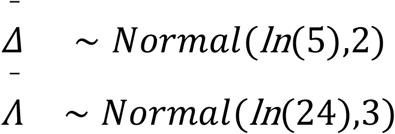

We used Normal priors centered on zero for the culture effects *X*_*n*_ and the surface effects *μ*_*n*_ with standard deviations designed to rule out implausibly large effects.

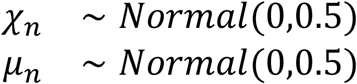

We placed Pos-Normal priors on the standard deviations *σ*_*Δ*_ and *σ*_*Λ*_:

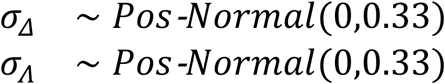

These allow for either substantial (1 log) variation compared to the regression prediction or for minimal variation.

All other priors were as in the half-life estimation model (section 3).

#### Predictive checks

We assessed the appropriateness of prior distribution choices using prior predictive checks and assessed goodness of fit for the estimated model using posterior predictive checks.

## ACKNOWLEDGMENTS

This work was funded by the National Institutes of Health (NIH) CEIRS (HHSN272201400007C) and New Innovator Award to LCM (1-DP2-A1112243), the American Lung Association Biomedical Research Grant (RG-575688), the Tsinghua Education Foundation of North America, and the China Scholarship Council (CSC). Work by DHM and JOL-S was supported by the UCLA AIDS Institute and Charity Treks, and National Science Foundation (NSF) Grants DEB-1557022 and DEB-2245631. We thank members of the Lakdawala Lab, Dr. Bill Goins (University of Pittsburgh), and the Marr lab at Virginia Tech for critical feedback during the studies, data analysis, and in manuscript preparation. We thank Dr. Rachel Duron for editorial assistance.

## REFERENCES

1. S. S. Shrestha et al., Estimating the burden of 2009 pandemic influenza A (H1N1) in the United States (April 2009-April 2010). Clin Infect Dis 52 Suppl 1, S75–82 (2011).

2. X. Xu et al., Update: Influenza Activity in the United States During the 2018–19 Season and Composition of the 2019–20 Influenza Vaccine. MMWR Morb Mortal Wkly Rep 68, 544–551 (2019).

3. G. Brankston, L. Gitterman, Z. Hirji, C. Lemieux, M. Gardam, Transmission of influenza A in human beings. Lancet Infect Dis 7, 257–265 (2007).

4. B. Killingley, J. Nguyen-Van-Tam, Routes of influenza transmission. Influenza Other Respir Viruses 7 Suppl 2, 42–51 (2013).

5. S. A. Boone, C. P. Gerba, Significance of fomites in the spread of respiratory and enteric viral disease. Appl Environ Microbiol 73, 1687–1696 (2007).

6. M. Nicas, R. M. Jones, Relative contributions of four exposure pathways to influenza infection risk. Risk Anal 29, 1292–1303 (2009).

7. N. Zhang, Y. Li, Transmission of Influenza A in a Student Office Based on Realistic Person-to-Person Contact and Surface Touch Behaviour. Int J Environ Res Public Health 15, (2018).

8. B. Bean et al., Survival of influenza viruses on environmental surfaces. J Infect Dis 146, 47–51 (1982).

9. S. A. Boone, C. P. Gerba, The occurrence of influenza A virus on household and day care center fomites. J Infect 51, 103–109 (2005).

10. A. D. Coulliette, K. A. Perry, J. R. Edwards, J. A. Noble-Wang, Persistence of the 2009 pandemic influenza A (H1N1) virus on N95 respirators. Appl Environ Microbiol 79, 2148–2155 (2013).

11. K. A. Perry et al., Persistence of Influenza A (H1N1) Virus on Stainless Steel Surfaces. Appl Environ Microbiol 82, 3239–3245 (2016).

12. K. A. Thompson, A. M. Bennett, Persistence of influenza on surfaces. J Hosp Infect 95, 194–199 (2017).

13. J. Zhao, J. E. Eisenberg, I. H. Spicknall, S. Li, J. S. Koopman, Model analysis of fomite mediated influenza transmission. PLoS One 7, e51984 (2012).

14. J. Guan, M. Chan, A. VanderZaag, Inactivation of Avian Influenza Viruses on Porous and Non-porous Surfaces is Enhanced by Elevating Absolute Humidity. Transbound Emerg Dis 64, 1254–1261 (2017).

15. J. Oxford et al., The survival of influenza A(H1N1)pdm09 virus on 4 household surfaces. Am J Infect Control 42, 423–425 (2014).

16. J. O. Noyce, H. Michels, C. W. Keevil, Inactivation of influenza A virus on copper versus stainless steel surfaces. Appl Environ Microbiol 73, 2748–2750 (2007).

17. S. L. Warnes, C. W. Keevil, Inactivation of norovirus on dry copper alloy surfaces. PLoS One 8, e75017 (2013).

18. S. L. Warnes, Z. R. Little, C. W. Keevil, Human Coronavirus 229E Remains Infectious on Common Touch Surface Materials. mBio 6, e01697–01615 (2015).

19. W. Yang, S. Elankumaran, L. C. Marr, Relationship between humidity and influenza A viability in droplets and implications for influenza’s seasonality. PLoS One 7, e46789 (2012).

20. J. H. Hemmes, K. C. Winkler, S. M. Kool, Virus survival as a seasonal factor in influenza and polimyelitis. Nature 188, 430–431 (1960).

21. L. C. Marr, J. W. Tang, J. Van Mullekom, S. S. Lakdawala, Mechanistic insights into the effect of humidity on airborne influenza virus survival, transmission and incidence. J R Soc Interface 16, 20180298 (2019).

22. J. W. Tang et al., Comparison of the incidence of influenza in relation to climate factors during 2000–2007 in five countries. Journal of medical virology 82, 1958–1965 (2010).

23. W. Yang, S. Elankumaran, L. C. Marr, Concentrations and size distributions of airborne influenza A viruses measured indoors at a health centre, a day-care centre and on aeroplanes. J R Soc Interface 8, 1176–1184 (2011).

24. L. Bourouiba, Fluid Dynamics of Respiratory Infectious Diseases. Annu Rev Biomed Eng 23, 547–577 (2021).

25. M. Moriyama, W. J. Hugentobler, A. Iwasaki, Seasonality of Respiratory Viral Infections. Annu Rev Virol 7, 83–101 (2020).

26. K. A. Kormuth et al., Influenza Virus Infectivity Is Retained in Aerosols and Droplets Independent of Relative Humidity. J Infect Dis 218, 739–747 (2018).

27. K. A. Kormuth et al., Environmental Persistence of Influenza Viruses Is Dependent upon Virus Type and Host Origin. mSphere 4, (2019).

28. Y. Thomas et al., Survival of influenza virus on banknotes. Appl Environ Microbiol 74, 3002–3007 (2008).

29. N. Pica, N. M. Bouvier, Environmental factors affecting the transmission of respiratory viruses. Curr Opin Virol 2, 90–95 (2012).

30. T. P. Weber, N. I. Stilianakis, Inactivation of influenza A viruses in the environment and modes of transmission: a critical review. J Infect 57, 361–373 (2008).

31. M. M. Myerburg, P. R. Harvey, E. M. Heidrich, J. M. Pilewski, M. B. Butterworth, Acute regulation of the epithelial sodium channel in airway epithelia by proteases and trafficking. Am J Respir Cell Mol Biol 43, 712–719 (2010).

32. C. Brownie et al., Estimating viral titres in solutions with low viral loads. Biologicals 39, 224–230 (2011).

33. D. H. Morris et al., Mechanistic theory predicts the effects of temperature and humidity on inactivation of SARS-CoV-2 and other enveloped viruses. Elife 10, (2021).

34. J. Han et al., Efficient and quick inactivation of SARS coronavirus and other microbes exposed to the surfaces of some metal catalysts. Biomed Environ Sci 18, 176–180 (2005).

35. A. L. Rasmussen, Probing the Viromic Frontiers. mBio 6, e01767–01715 (2015).

36. J. D. Recker, X. Li, Evaluation of Copper Alloy Surfaces for Inactivation of Tulane Virus and Human Noroviruses. J Food Prot 83, 1782–1788 (2020).

37. N. van Doremalen et al., Aerosol and Surface Stability of SARS-CoV-2 as Compared with SARS-CoV-1. N Engl J Med 382, 1564–1567 (2020).

38. H. T. Michels, J. O. Noyce, C. W. Keevil, Effects of temperature and humidity on the efficacy of methicillin-resistant Staphylococcus aureus challenged antimicrobial materials containing silver and copper. Lett Appl Microbiol 49, 191–195 (2009).

39. S. L. Warnes, C. W. Keevil, Lack of Involvement of Fenton Chemistry in Death of Methicillin-Resistant and Methicillin-Sensitive Strains of Staphylococcus aureus and Destruction of Their Genomes on Wet or Dry Copper Alloy Surfaces. Appl Environ Microbiol 82, 2132–2136 (2016).

40. L. Weaver, J. O. Noyce, H. T. Michels, C. W. Keevil, Potential action of copper surfaces on meticillin-resistant Staphylococcus aureus. J Appl Microbiol 109, 2200–2205 (2010).

41. G. Borkow, S. S. Zhou, T. Page, J. Gabbay, A novel anti-influenza copper oxide containing respiratory face mask. PLoS One 5, e11295 (2010).

42. A. A. Cortes, J. M. Zuniga, The use of copper to help prevent transmission of SARS-coronavirus and influenza viruses. A general review. Diagn Microbiol Infect Dis 98, 115176 (2020).

43. Y. Fujimori et al., Novel antiviral characteristics of nanosized copper(I) iodide particles showing inactivation activity against 2009 pandemic H1N1 influenza virus. Appl Environ Microbiol 78, 951–955 (2012).

44. D. A. Edwards et al., Exhaled aerosol increases with COVID-19 infection, age, and obesity. Proc Natl Acad Sci U S A 118, (2021).

45. D. C. Adam et al., Clustering and superspreading potential of SARS-CoV-2 infections in Hong Kong. Nat Med 26, 1714–1719 (2020).

46. S. S. Lakdawala, V. D. Menachery, Catch Me if You Can: Superspreading of COVID-19. Trends Microbiol 29, 919–929 (2021).

47. J. O. Lloyd-Smith, S. J. Schreiber, P. E. Kopp, W. M. Getz, Superspreading and the effect of individual variation on disease emergence. Nature 438, 355–359 (2005).

48. L. C. Marr, J. W. Tang, A Paradigm Shift to Align Transmission Routes With Mechanisms. Clin Infect Dis 73, 1747–1749 (2021).

49. L. J. Reed, H. Muench, A simple method of estimating fifty percent endpoints. Am J Hyg 27, 493–497 (1938).

